# A gnotobiotic system reveals multifunctional effector roles in plant-fungal pathogen dynamics

**DOI:** 10.1101/2025.03.27.645772

**Authors:** Wilko Punt, Jiyeun Park, Hanna Roevenich, Anton Kraege, Natalie Schmitz, Jan Wieneke, Nick C. Snelders, Gabriel L. Fiorin, Ana López-Moral, Edgar A. Chavarro-Carrero, Gabriella C. Petti, Kathrin Wippel, Bart P.H.J. Thomma

**Affiliations:** University of Cologne, Institute for Plant Sciences, Cluster of Excellence on Plant Sciences (CEPLAS), 50674 Cologne, Germany; Wageningen University and Research, Laboratory of Phytopathology, 6708PB Wageningen, The Netherlands; University of Amsterdam, Swammerdam Institute for Life Sciences, 1000BE Amsterdam, The Netherlands

## Abstract

Plants host diverse microbiota that influence physiological processes and can enhance resilience against invading pathogens that, in turn, evolved effector proteins to manipulate host microbiota in their favor. However, the complexity of microbial communities and their interactions complicates mechanistic research on processes governing microbiota assembly and function. Gnotobiotic systems are valuable tools to study plant microbiota by reducing complexity and enabling controlled microbiota reconstitution experiments. Despite their utility, no gnotobiotic systems have been established to investigate the role of antimicrobial effector proteins in the interactions between plants, their microbiota, and fungal pathogens. Here, we present a refined gnotobiotic system designed to study these interactions, establishing protocols for infections with the fungal pathogen *Verticillium dahliae* across multiple host plants under sterile conditions. We demonstrate that a synthetic microbial community (SynCom) derived from a culture collection generated for this study can be applied in this system where it interferes with fungal infections. Additionally, using our gnotobiotic system we reveal that specific antimicrobial effectors of *V. dahliae,* like Ave1L2, contribute to fungal virulence in a microbiota-dependent manner, whereas other antimicrobial effectors, such as Ave1, seem to possess functions beyond microbiota manipulation.

## Introduction

During their life, plants functions as holobionts, an integrated unit consisting of the plant with its associated microbial communities, also known as the plant microbiota (Vandenkoornhuyse *et al*. 2015). This complex microbiota is associated with all plant parts and its composition depends on the interplay of various biotic and abiotic factors (Trivedi *et al*. 2020; Mesny et al., 2023). Microbes that establish in the plant microbiota can interact with the plant host in various ways, ranging from beneficial and growth-promoting to pathogenic and disease-inducing (Mesny *et al*. 2024). Consequently, the composition and functions of the microbiota contribute to plant health and productivity. Importantly, plants possess the ability to actively shape their microbiota to alleviate biotic or abiotic stresses. For instance, upon invasion of the “take-all-disease” pathogen *Gaeumannomyces graminis* var. *tritici*, wheat plants in particular fields in the USA were demonstrated to actively recruit beneficial *Pseudomonas* bacteria into their microbiota. These *Pseudomonas* species antagonize *G. graminis* via the secretion of 2,4-diacetylphloroglucinol or phenanzine-1-carboxylic acid antibiotics, leading to “take-all decline”; a reduction of disease severity over time (Spooren *et al*. 2024). Thus, microbiota have been described as an additional layer of plant defense against pathogen invasion (Mendes *et al*. 2011; Carrión *et al*. 2019). Microbiota not only serve as crucial defense barrier against invading pathogens, but also help plants mitigate the pathogenic potential of certain microbial community members. This is evident in cases where some microbes severely inhibit plant growth when inoculated individually, yet this negative impact disappears when they are introduced within a community context (Durán *et al*. 2018).

During host invasion, plant pathogens secrete so-called effectors, a diversity of molecules some of which remain in the apoplast while others have intracellular destinations where they function to promote host colonization (Jones and Dangl 2006; Cook *et al*. 2015). Initially, effectors were proposed to interfere with host immune responses, by interfering with pathogen perception by plant immune receptors, downstream immune signalling, or the execution of immunity (de Jonge *et al*. 2010; Bozkurt *et al*. 2011; King *et al*. 2014). However, later it was realized that besides direct modulation of plant immune responses, effectors can also function in self-protection to undermine plant immune responses in an indirect manner (van den Burg *et al*. 2006; de Jonge *et al*. 2010). Although effectors were initially implicated in suppression of host immune responses, manipulation of other host physiological processes could also be demonstrated, including for instance the induction of water-soaking by manipulation of stomatal closure, or sugar release into the apoplast through the manipulation of host sugar transporter expression (Chen *et al*. 2010; Hu *et al*. 2022; Roussin-Léveillée *et al*. 2022). Notably, several effectors were described to possess multiple functions, contributing to host colonization through different mechanisms (Liu *et al*. 2016; Lin *et al*. 2023).

Recently it was shown that pathogens exploit effector proteins to target host microbiota, and manipulate their composition to breach the protective defense layer that the microbiota provides which ultimately promotes host colonization (Kettles *et al*. 2018; Snelders *et al*. 2020; Snelders *et al*. 2021; Chang *et al*. 2021; Snelders *et al*. 2023; Ökmen *et al*. 2023; Gómez-Pérez *et al*. 2023; Chavarro-Carrero *et al*. 2024; Mesny *et al*. 2024). For example, the effector protein Ave1 from the fungal plant pathogen *Verticillium dahliae* exhibits selective antibacterial activity, suppressing Sphingomonadales bacteria in the rhizosphere microbiota of infected cotton and tomato plants. Since Sphingomonadales can reduce Verticillium wilt severity, *V. dahliae* exploits Ave1 to inhibit these beneficial bacteria and facilitate host colonization (Snelders *et al*. 2020). Building on this discovery, the effector protein Ave1L2 was identified through sequence similarity to Ave1, and similarly exhibits selective antibacterial activity, albeit with a distinct activity spectrum. *In planta* assays showed that Ave1L2 manipulates host microbiota by targeting Actinobacteria in the tomato rhizosphere. Antibiotic-induced depletion of Actinobacteria in the plant microbiota increased plant susceptibility to *V. dahliae* while reducing the virulence impact of Ave1L2 (Snelders *et al*. 2023). In both cases, the contribution of antimicrobial effector proteins to virulence of *V. dahliae* could only be tested in the presence of plant-associated microbiota, making it impossible to exclude the simultaneous occurrence of virulence contributions through direct manipulation of host targets. Considering that several antimicrobial effectors seem to have an ancient origin, and likely acted in intermicrobial competition before land plant evolution, they may have acquired additional functions in host manipulation during fungal co-evolution with their host plants (Snelders *et al*. 2021; Snelders *et al*. 2022; Mesny and Thomma 2024).

Research on plant-microbiota interactions, including the role and contribution of antimicrobial effector proteins, is often complicated by the sheer complexity of host-associated microbial communities. The numerous plant-microbe and intermicrobial interactions, which constantly respond to various environmental cues, make mechanistic studies challenging. Gnotobiotic plant growth systems, which allow for controlled experiments in the presence or absence of particular microbiota, offer a powerful tool to address these challenges (Vorholt *et al*. 2017; Liu *et al*. 2019; Ma *et al*. 2022). Over the past decades, various gnotobiotic plant growth systems have been developed based on diverse substrates, including agar-based (Innerebner *et al*. 2011), clay-based (Carlström *et al*. 2019), and peat-based substrates (Kremer *et al*. 2021), each with distinct advantages and limitations. Agar-based systems provide precise control over nutrient availability but generate highly artificial conditions. Clay-based systems offer a soil-like substrate structure and are easily sterilizable, but lack organic carbon, and the substrate itself makes it challenging to regulate nitrogen and phosphorus levels. In contrast, peat-based systems comprise organic carbon, yet lack control over nutrient composition (Liu *et al*. 2019; Kremer *et al*. 2021). Ultimately, the choice of gnotobiotic system depends on the specific research question that is addressed. Gnotobiotic systems are frequently used in reductionist experiments and can be particularly powerful when combined with synthetic microbial communities (SynComs). Such SynComs typically are communities with reduced complexity when compared with natural communities, generated from microbial culture collections, allowing to monitor the impact of defined microbial communities on a particular trait of interest (Vorholt *et al*. 2017; Novak *et al*., 2024). For example, experiments utilizing a calcined clay-based gnotobiotic system and a culture collection of *Arabidopsis thaliana*-associated bacteria were used to demonstrate the role of priority effects during assembly of the *A. thaliana* phyllosphere microbiota (Carlström *et al*. 2019). Additionally, a repopulation study using *A. thaliana* plants grown in a peat-based gnotobiotic system with a 106-member multi-kingdom SynCom identified evolutionary conserved genetic determinants for bacterial root colonization (Vannier *et al*. 2023). Several studies have utilized gnotobiotic systems to study the impact of the microbiota on plant defense against pathogens (Vogel *et al*. 2021; Paasch *et al*. 2023). For instance, in a systematic approach, screening of 224 bacterial isolates from an *A. thaliana* phyllosphere culture collection for protection against the bacterial pathogen *Pseudomonas syringe* revealed that 10% of the bacteria can prevent bacterial speck disease (Vogel *et al*. 2021).

Due to the lack of suitable inoculation protocols, no gnotobiotic system has been available to determine whether antimicrobial effector proteins from the soil-borne fungal pathogen *V. dahliae* contribute to virulence also in the absence of host-associated microbiota, and therefore have additional host targets. In this study, we describe a peat-based gnotobiotic system for plant inoculations with *V. dahliae* and its use to investigate the role of antimicrobial effector proteins in fungal virulence.

## Results

### Establishment of a Flowpot-system tailored for Verticillium wilt development

We aimed to modify a previously published Flowpot-system to study the behavior of the fungal pathogen *Verticillium dahliae* during disease development (Kremer *et al*. 2021). Our adapted Flowpot-system relies on plants grown in commonly available 50 ml syringes that are filled with a blend of peat substrate and vermiculite (Figure 1a). The substrate mixture is sterilized in three consecutive autoclaving steps. To assess sterility, sterilized substrate was plated onto various growth media. Following four days of incubation, no signs of microbial growth were observed, indicating successful substrate sterilization (Suppl. Figure 1a). Next, we aimed to generate a non-sterile control substrate that underwent a similar treatment through triple sterilization. For this purpose, we recolonized sterilized substrate by mixing sterilized and non-sterilized substrate in a 9:1 ratio (Figure 1b). To verify substrate recolonization, we again made use of plating, revealing substantial microbial growth on all media (Suppl. Figure 1a).

**Figure 1.**
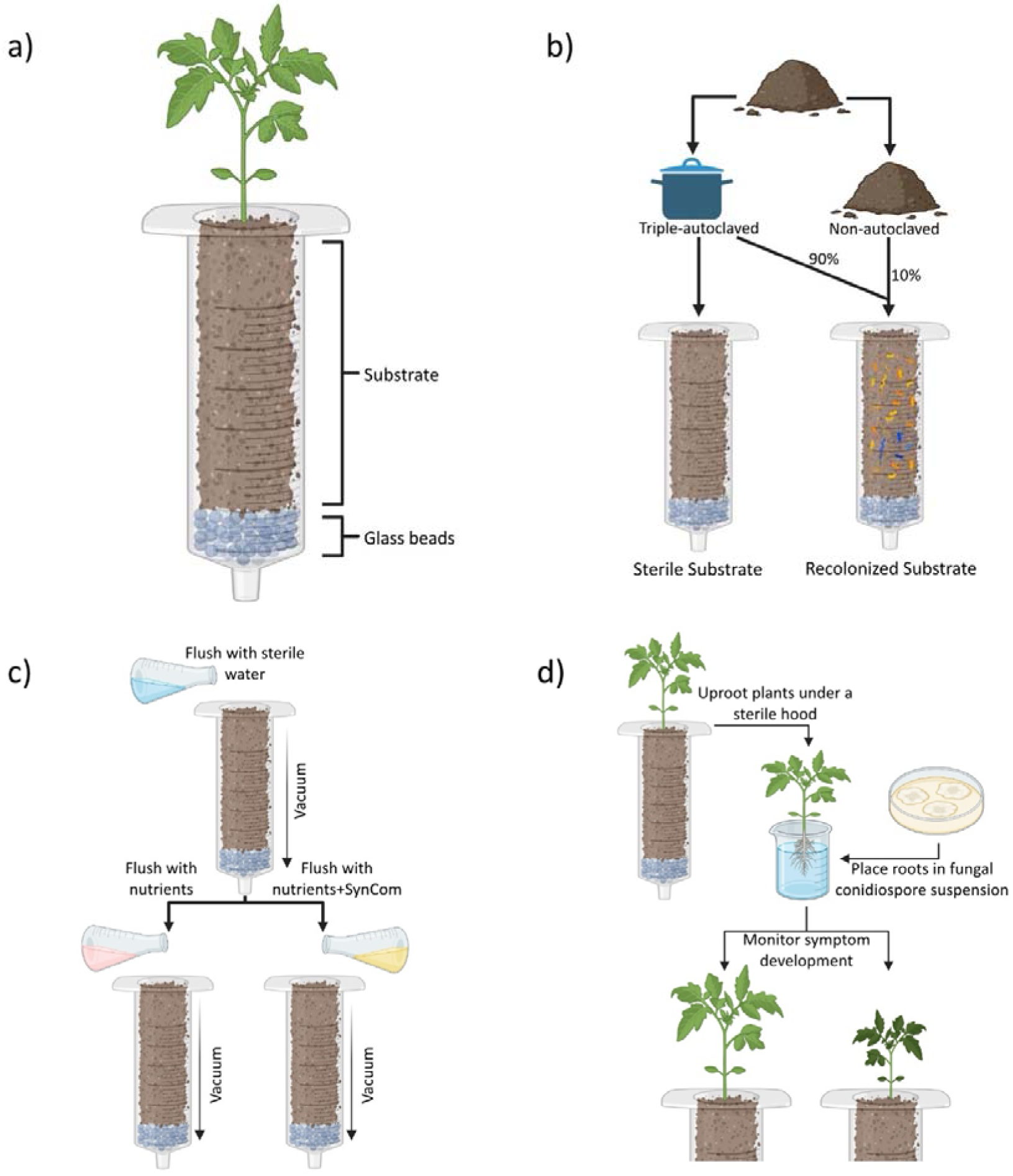
Technical set-up of the Flowpot-ystem. **a)** Schematic overview of an individual Flowpot-unit. **b)** Substrate preparation procedure. Triple autoclaved substrate is mixed with 10% untreated soil to create a substrate that is recolonized by microbes. **c)** Flowpot-flushing procedure. Water is flushed through each Flowpot using vacuum, followed by another flushing step with a nutrient solution that can be supplemented with a SynCom. **d)** *Verticillium dahliae* inoculation procedure. Plants are uprooted in a sterile hood and placed into V. dahliae conidiospore-suspension for several minutes before replanting into the substrate and monitored for symptom development at 14 dpi.

Considering the impact of autoclaving on the physicochemical properties of peat-based substrates and to try and eliminate potentially toxic compounds that might have been released during the autoclaving procedure that may affect plant growth, we applied substrate washes. To this end, vacuum was applied to each Flowpot and the substrate was rinsed with sterilized water. Moreover, the flushing mechanism was utilized to supply nutrients to the substrate (Figure 1c).

Gnotobiotic systems can be used for microbiota reconstitution experiments, conducted through the application of single microbial species or SynComs to study the role of defined microbiota on a particular trait. To enable inoculation of the substrate, our Flowpot protocol utilizes the vacuum flushing mechanism to supplement substrate with microbial suspensions, thereby enabling substrate colonization by a defined microbial inoculum. In this manner, our modified Flowpot-system provides a versatile platform for experiments to be conducted either in the absence or in the presence of non-defined or defined microbial communities.

### The Flowpot-system is suitable for *Verticillium* infections on various host plants

Thus far, the role and impact of antimicrobial effector proteins of the fungal plant pathogen *V. dahliae* has been studied mostly on tomato. Research on these proteins can be facilitated substantially with suitable gnotobiotic systems, allowing for fungal infections in otherwise sterile environments. Therefore, we aimed to establish infections of *V. dahliae* on tomato plants in our Flowpot-system. To test if the Flowpot-system is suitable to maintain tomato plants under gnotobiotic conditions, we germinated surface-sterilized seeds on sterilized substrate and assessed plant growth. Despite a low germination rate of only 20% under these conditions, tomato plants grown in gnotobiotic conditions appeared healthy after 24 days of growing, indicating that the conditions are suitable for growth (Figure 2a). Since plants recruit a substantial part of their microbiota from the surrounding bulk soil, but also from endophytes that already reside within the plant seed, we also assessed to what extent growth in the sterilized substrate leads to a less diverse plant microbiota. To this end, we conducted 16S amplicon sequencing of tomato stems of plants grown on sterilized or recolonized substrate. As expected, plants grown on sterilized substrate carried communities with a significantly lower Shannon index when compared with plants grown on recolonized substrate, indicating that plants grown on sterilized substrate harbor significantly less diverse microbial communities (Suppl. Figure 1b). This reduction in microbial diversity correlated with notable effects on plant growth, as tomato plants grown on recolonized substrate revealed substantially higher germination rates (76%) and produced significantly more biomass when compared with plants grown on sterilized substrate, demonstrating the growth-promoting ability of diverse microbiota (Figure 2a).

**Figure 2.**
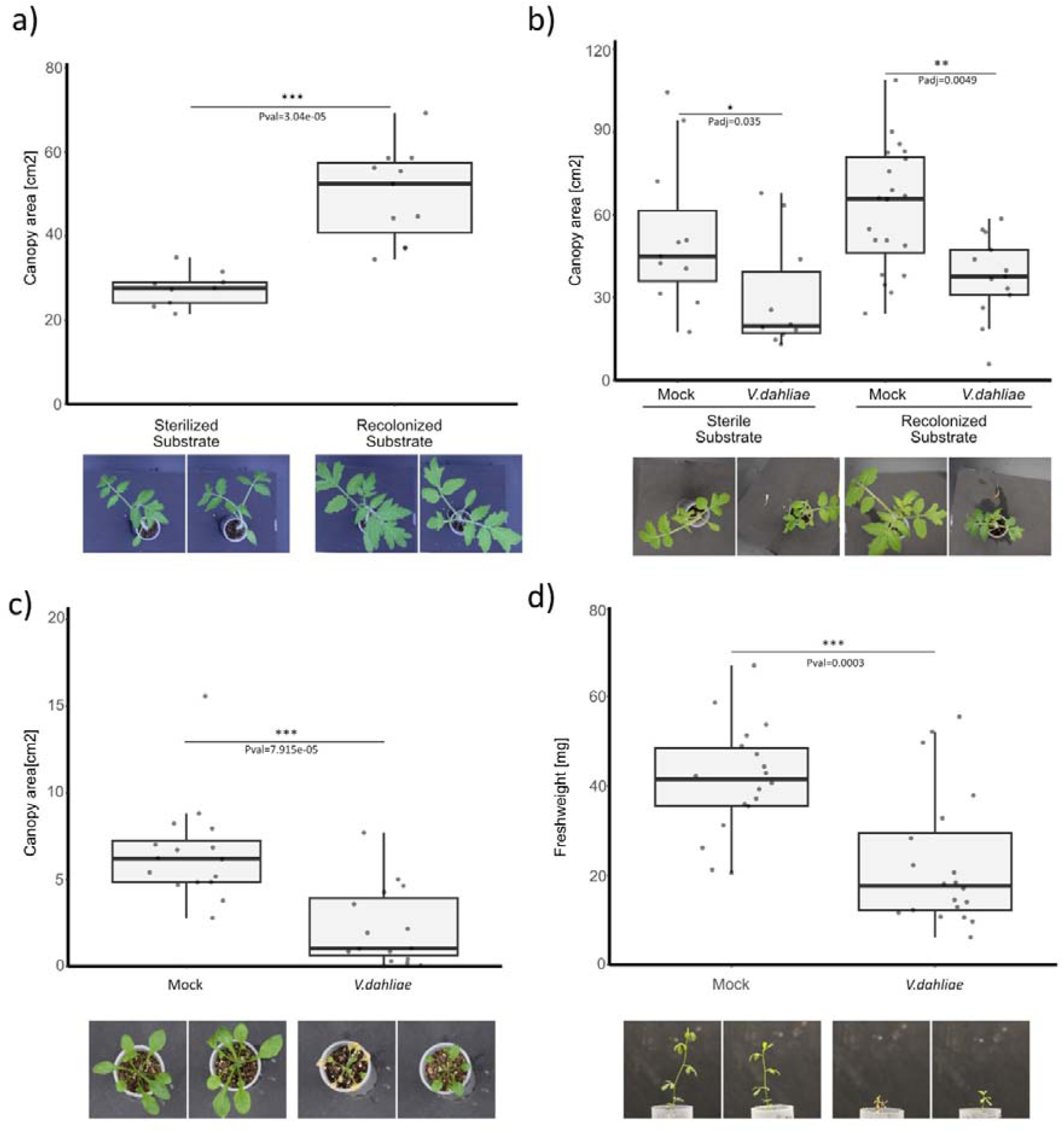
The Flowpot-system accommodates interactions of *V. dahliae* with diverse host plants. **a)** Canopy area of tomato plants grown on either sterilized or recolonized substrate. Pairwise comparison was performed using the Wilcoxon Rank Sum test (*** = Pval < 0.001). **b)** Canopy area of tomato plants grown on recolonized or sterilized substrate at 14 dpi with *V. dahliae*. Pairwise comparison was performed using the Wilcoxon Rank-Sum test, followed by multiple testing correction using Benjamini-Hochberg (FDR) corrections. (* = Padj < 0.05; **= Padj < 0.01). **c)** Canopy area of *Arabidopsis thaliana*Col-0 plants grown in sterilized substrate at 14 dpi of *V. dahliae*. Pairwise comparison was performed using the Wilcoxon Rank Sum test (*** = Pval < 0.001). **d)** Fresh weight of *Lotus japonicus* cv. Gifu plants grown on sterilized substrate at 14 dpi of *V. dahliae*. Pairwise comparison was performed using the Wilcoxon Rank Sum test (*** = Pval < 0.001).

Next, we aimed to establish *V. dahliae* infections on tomato plants grown in our gnotobiotic system. To this end, tomato seedlings grown on either sterilized or recolonized substrate were uprooted in a sterile hood, inoculated with a *V. dahliae* conidiospore suspension, and replanted in the Flowpots (Figure 1d). At 14 days post inoculation, *V. dahliae-*inoculated plants grown in recolonized substrate displayed significantly less growth when compared with mock-inoculated plants. Similarly, also in sterilized substrate, plants treated with *V. dahliae* revealed significantly reduced growth when compared with mock-inoculated plants, indicating successful *V. dahliae* infection under the gnotobiotic conditions (Figure 2b). Considering the broad host range of *V. dahliae*, we investigated whether our inoculation protocol is also suitable for other plant hosts. To this end, we tested *Arabidopsis thaliana*and *Lotus japonicus*, as both plant species were previously grown in the Flowpot-system (Kremer *et al*. 2021; Wippel *et al*. 2021). Interestingly, both *A. thaliana* and *L. japonicus* plants displayed significantly reduced growth upon *V. dahliae* inoculation when compared with mock-inoculated plants, indicating that our *V. dahliae* inoculation protocol is suitable to study diverse *V. dahliae*-host interactions under gnotobiotic conditions (Figure 2c, d).

### Application of a protective SynCom prevents Verticillium wilt symptoms

To enable reconstitution experiments on tomato plants using host-associated bacteria, we generated a bacterial culture collection of tomato-associated bacteria. To this end, two commercially purchased tomato plants were separated into phyllosphere and root samples, which served as starting material for a colony picking approach. In total, 374 colonies were picked and identified using Sanger sequencing followed by BLAST searches to the NCBI database, which led to a total number of 132 unique bacterial isolates. Further, we confirmed the results from the BLAST identification by performing whole-genome sequencing on 75 of the species with Oxford Nanopore Technology (ONT) Sequencing. The culture collection comprises 100 distinct isolates that belong to 48 genera isolated from root tissue, and 48 distinct isolates that belong to 31 genera isolated from phyllosphere tissues (Figure 3a).

**Figure 3.**
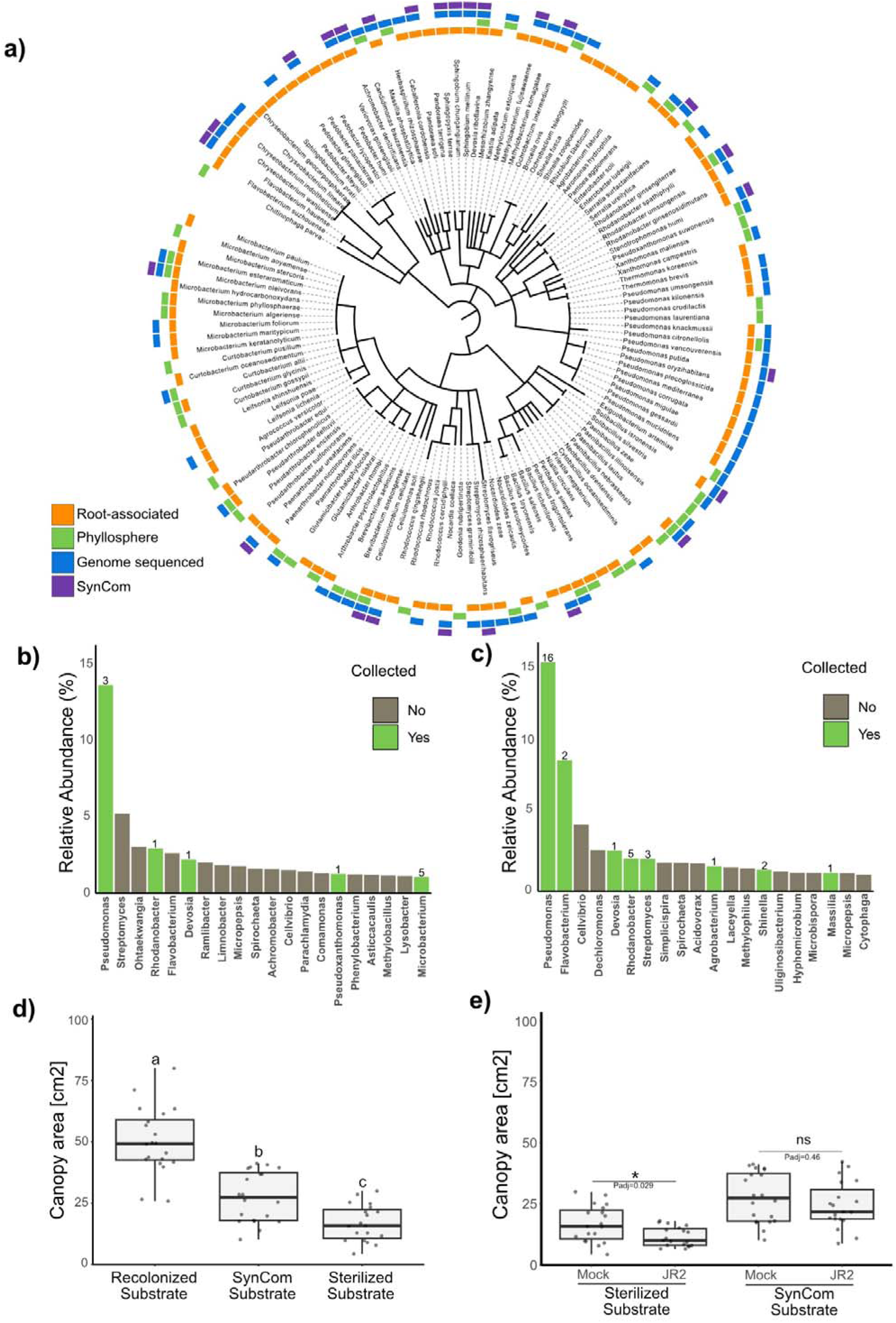
Tomato-derived synthetic communities can suppress Verticillium wilt disease. **a)** Phylogenetic tree of the tomato-associated culture collection. Orange boxes indicate strains isolated from the rhizosphere, while green boxes indicate strains isolated from the phyllosphere. Blue boxes indicate strains that were subjected to whole genome sequencing. Purple boxes indicate strains that were used to compose the SynCom. **b)** Relative abundance in % of the 20 most abundant genera in the tomato phyllosphere microbiota. Green bars indicate genera of which at least one member was isolated, whereas grey bars indicate genera of which no member was isolated. Numbers above the bars indicate the number of strains isolated per genus. **c)** Relative abundance in % of the 20 most abundant genera in the tomato rhizosphere microbiota. Green bars indicate genera of which at least one member was isolated, whereas grey bars indicate genera of which no member was isolated. Numbers above the bars indicate the number of strains isolated per genus. **d)** Canopy area of plants grown on either sterilized, SynCom-treated or recolonized substrate. Different letters indicate statistical differences based on One-Way-Anova (Tukey HSD-Test pval < 0.05). **e)** Canopy area of mock- or *V. dahliae*-inoculated plants grown on sterilized or SynCom-treated substrate. Pairwise comparisons are performed using Wilcoxon Rank-Sum test, followed by multiple testing correction using Benjamini-Hochberg (FDR) correction

Next, we compared the species isolated by our colony picking approach to the microbiota of the input tomato material. To this end, we conducted 16S amplicon sequencing on the plant material and determined the most abundant bacterial genera. The 20 most abundant genera compose 55% of the tomato root microbiota, with *Pseudomonas*, *Flavobacterium* and *Cellvibrio* as the most abundant genera. In the tomato phyllosphere microbiota, the 20 most abundant genera make up 50% of the input microbiota with *Pseudomonas*, *Streptomyces* and *Ohtaekwangia* as most abundant. The root-associated culture collection contains at least one isolate from eight of the 20 most abundant genera, with 74 isolates belonging to less abundant genera. The phyllosphere culture collection captured at least one isolate from 5 out of the 20 most abundant genera, with 37 isolates belonging to other, less abundant genera (Figure 3b). Thus, our culture collection captured a wide diversity of microbes from the tomato microbiota.

To employ this culture collection within our Flowpot-system, we generated a SynCom composed of 26 isolates in an attempt to reduce the impact of *V. dahliae* infection on tomato plants. The selection of isolates was focused on the root-associated collection, to enhance the likelihood of successful substrate colonization. We selected one representative isolate from each family that is present in the root-associated collection and prioritized the selection of species that were previously reported to suppress disease or promote growth. First, we assessed if the application of the SynCom leads to rescue of the growth depletion phenotype we observed for plants grown in sterilized substrate. To this end, we grew plants on sterilized, recolonized and on sterilized substrate that was treated with the SynCom, respectively. Although tomato growth on SynCom-treated substrate was significantly reduced when compared with growth on recolonized substrate, tomato plants grown on SynCom-treated substrate produced significantly more biomass than plants grown on sterile substrate, suggesting that the SynCom partially restored the growth reduction observed in the absence of a substrate microbiota (Figure 3d). Next, we tested if pre-treatment of the substrate with the SynCom represses *V. dahliae* symptom development. Whereas plants grown on sterilized substrate revealed significant stunting upon *V. dahliae* inoculation in absence of the SynCom treatment, SynCom-treatment of the substrate eliminated *V. dahliae* symptom development, demonstrating that the SynCom successfully prevented disease development (Figure 3e).

### Discrimination of microbiota modulation from host target manipulation

Antimicrobial effector proteins of *V. dahliae* contribute to fungal virulence. However, due to the lack of suitable gnotobiotic systems it remained elusive if this virulence contribution is solely through microbiota manipulation, or rather relies on additional activities on plant virulence targets. If these effectors primarily function to manipulate host-associated microbiota, no contribution to fungal virulence should occur in plants that are devoid of an associated microbiota. To address this hypothesis, we grew tomato plants on sterilized or recolonized substrates and inoculated with *V. dahliae* wild-type strains or a deletion mutant for the gene encoding the antimicrobial effector protein Ave1L2. We previously showed that *V. dahliae* secretes Ave1L2 to facilitate host colonization of tomato plants by suppression of Actinobacteria. Reducing the abundance of Actinobacteria in the plant microbiota through antibiotic application led to increased plant sensitivity to *V. dahliae*, while reducing the virulence contribution of Ave1L2, suggesting that the effector is secreted to target plant-protective Actinobacteria (Snelders *et al*. 2023). In the Flowpot-system, plants that were grown in sterilized substrate revealed no difference in disease development when inoculated with wild-type *V. dahliae* or the Ave1L2 deletion mutant. In contrast, when grown on recolonized substrate, plants inoculated with the *Ave1L2* deletion mutant showed significantly reduced disease development when compared with plants that were inoculated with wild-type *V. dahliae*. Thus, Ave1L2 only contributes to virulence in the presence of a plant-associated microbiota, suggesting that this effector does not have additional virulence targets in the plant (Figure 4a).

**Figure 4.**
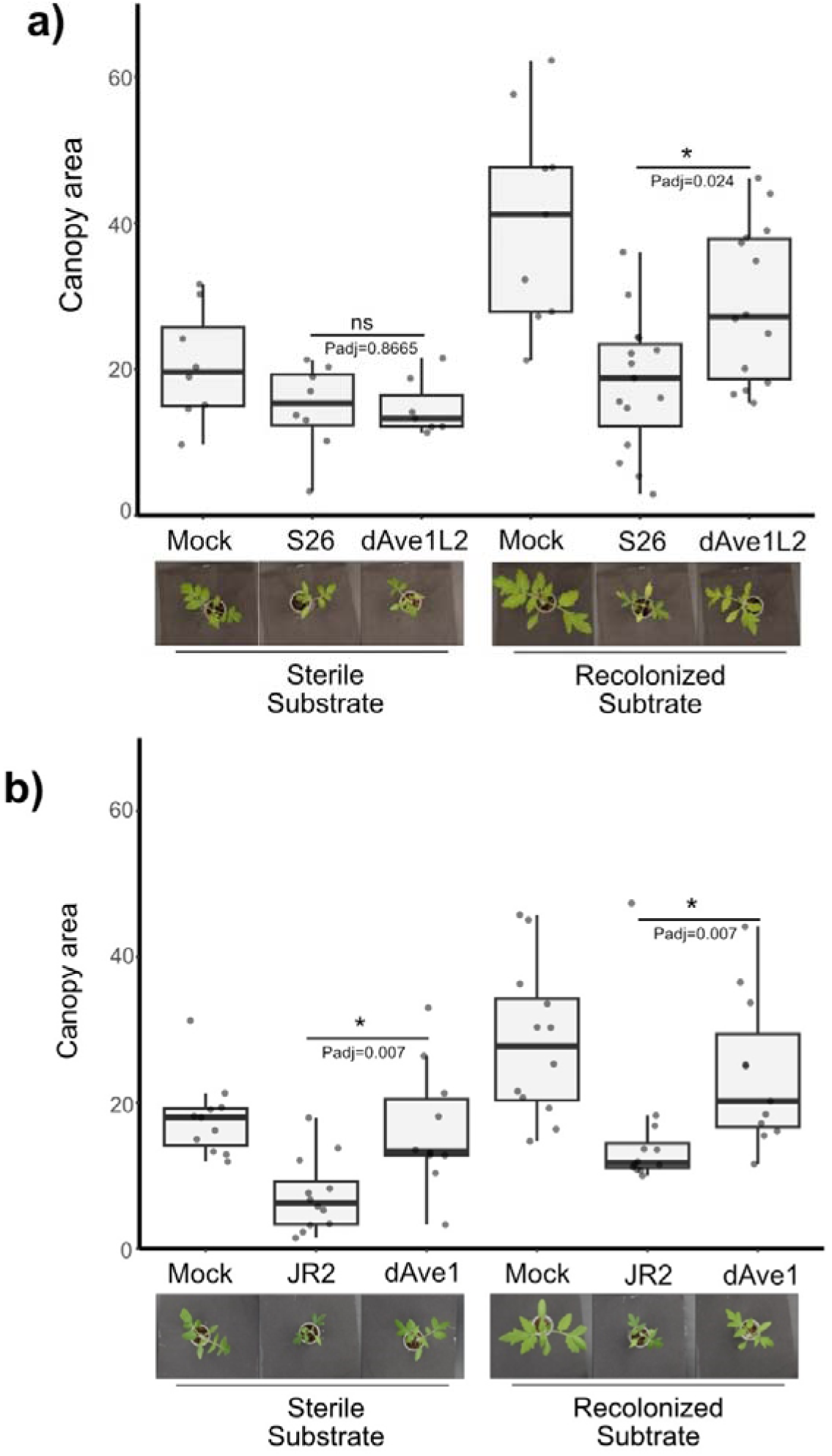
Antimicrobial effectors of *V. dahliae* differentially contribute to virulence in gnotobiotic conditions. **a)** Canopy area of tomato plants grown on either sterilized or recolonized substrate at 14 dpi of wild-type *V. dahliae* or an *Ave1L2* deletion mutant. Pairwise comparisons are performed using the Wilcoxon Rank-Sum test, followed by multiple testing correction using Benjamini-Hochberg (FDR) correction. **b)** Canopy area of tomato plants grown on either sterilized and recolonized substrate at 14 dpi of wild-type *V. dahliae* or an *Ave1* deletion mutant. Pairwise comparisons are performed using Wilcoxon Rank-Sum test, followed by multiple testing correction using Benjamini-Hochberg (FDR) correction.

Next, we assessed the virulence contribution of the *V. dahliae* Ave1 effector in sterilized substrate. This effector protein is utilized by *V. dahliae* to facilitate host colonization of tomato and cotton plants by suppression of Sphingomonad bacteria. We previously showed that pre-treatment of surface-sterilized tomato seeds with Sphingomonad bacteria reduced Verticillium wilt disease development, and that Ave1 secretion by *V. dahliae* significantly reduced Sphingomonad proliferation *in planta* (Snelders *et al*. 2020). In contrast to our observations for Ave1L2, tomato plants that were grown in sterilized substrate and inoculated with the *V. dahliae* wild-type strain JR2 were more stunted when compared with plants that were inoculated with an *Ave1* deletion mutant, indicating a clear virulence contribution of the effector protein on plants grown in sterile substrate. This contribution was also observed on plants grown on recolonized substrate, overall indicating that Ave1 may contribute to virulence in the absence of a host-associated microbiota too, which may rely on modulation of a host virulence target (Figure 4b).

### Ave1 affects host physiology

The *V. dahliae* effector Ave1 has numerous homologs in plants and in several other microbes, including *A. thaliana* PNP AtPNP-A and *Xanthomonas citri* pv. *citri* PNP XacPNP (de Jonge *et al*. 2012). Most plant homologs of Ave1 are annotated as plant natriuretic peptides (PNPs), mobile molecules that are released under biotic and abiotic stress conditions and have been implicated in several responses important for plant growth and homeostasis (Gehring and Irving 2003; Ruzvidzo *et al*. 2011). Multiple sequence alignment of the protein domains that have previously been implicated in PNP activity of AtPNP-A and XacPNP with Ave1 revealed high sequence similarity at the PNP site, a 12 amino acid long stretch that was previously reported to confer biological activity (Gottig *et al*., 2008). Notably, this similarity was much lower for Ave1L2 (Figure 5a). Previously PNPs were shown to be able to induce stomatal opening (Gottig *et al*. 2008). To investigate if Ave1 also possess such PNP-activity, we measured its ability to promote stomatal opening. Treatment of tomato leaf epidermis with purified Ave1 resulted in significantly enhanced stomatal opening as observed upon treatment with XacPNP and the synthetic auxin analogue indole-3-acetic acid (IAA) (Figure 5b). In contrast, addition of Ave1L2 did not affect stomatal opening. Consistent with previous reports demonstrating that PNP-induced responses are dependent on cyclic guanosine monophosphate (cGMP) signaling (Pharmawati *et al*. 2001; Turek and Gehring 2016), aperture changes caused by Ave1 were partially repressed by the guanylate cyclase inhibitor methylene blue (Supplementary Figure 2). Collectively, these findings demonstrate that Ave1 possess PNP activity, suggesting that beyond its antimicrobial function, Ave1 may also contribute to virulence through an additional function that involves the manipulation of host physiology.

**Figure 5.**
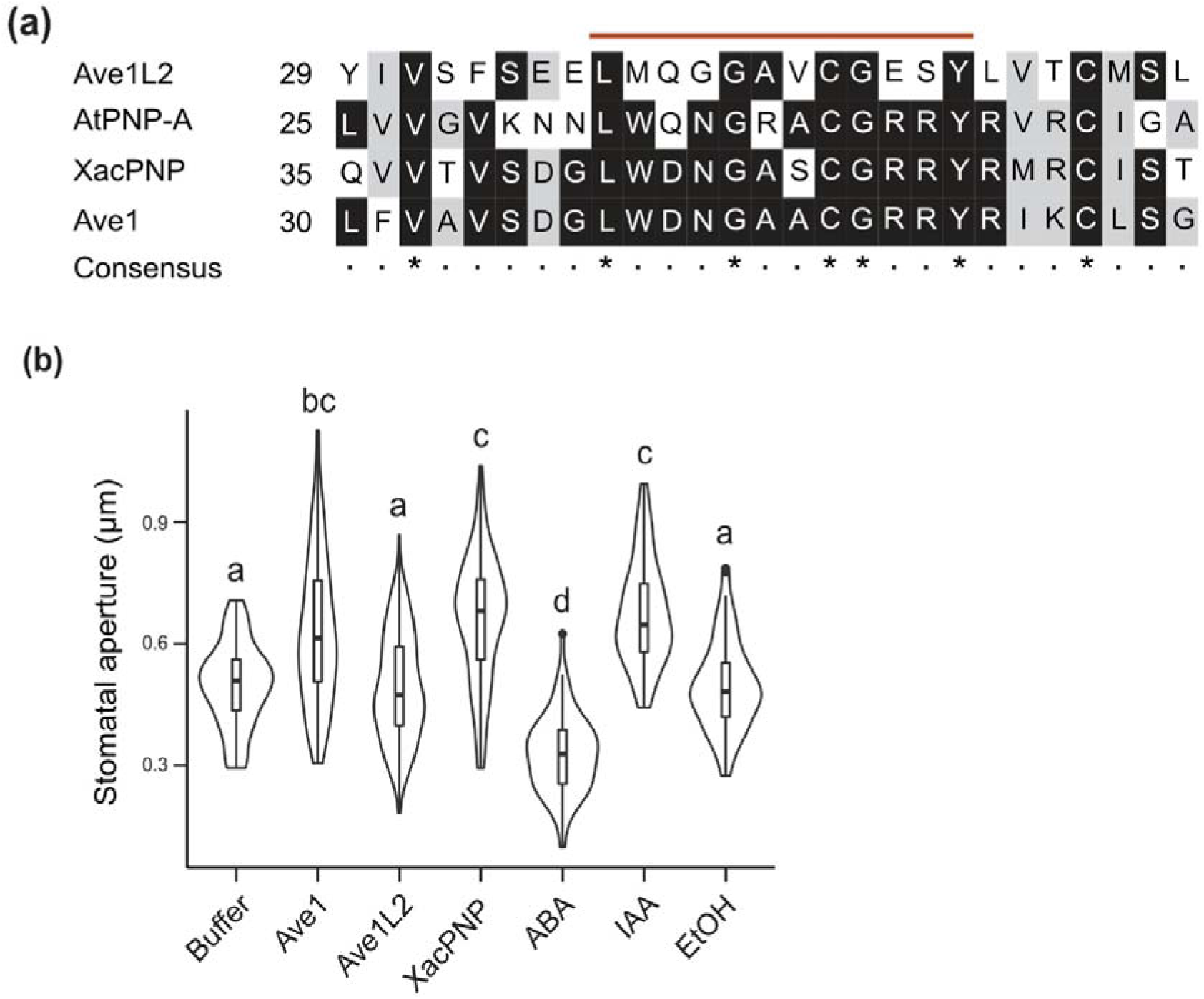
The *Verticillium dahliae* effector Ave1 contains an active PNP site, while Ave1L2 does not. **(a)**, Multiple sequence alignment of a short peptide stretch of Ave1 (amino acid 30 to 56) containing the PNP site (red line) with Ave1L2 and homologous sequences from *Xanthomonas citri* pv. *citri* (Xac) and *Arabidopsis thaliana*(At). (b), Stomatal opening in tomato epidermis following treatment with 5 µM purified protein. Indole-3-acetic acid (IAA; 1 µM) was used as positive control, whereas 50 µM abscisic acid (ABA) and EtOH served as negative controls. Data are from one representative experiment are shown. Letters represent statistically significant differences in stomatal opening index (width/length) according to one-way ANOVA (F (7,626) = 47.06, p<2e-16) and Tukey’s post-hoc test. Error bars represent the mean ± SD (n>60).

## Discussion

Over the past decades, research has established that plant-associated microbiota play crucial roles in plant health by suppressing the pathogenic potential of resident microbes and forming a barrier against invading pathogens (Mesny *et al*. 2023; Trivedi *et al*. 2020; Mesny *et al*. 2024). However, pathogens have evolved to overcome this additional layer of defense by secreting effector proteins with antimicrobial properties that manipulate host microbiota to their advantage (Kettles *et al*. 2018; Snelders *et al*. 2020; Snelders *et al*. 2021; Chang *et al*. 2021; Snelders *et al*. 2023; Ökmen *et al*. 2023; Gómez-Pérez *et al*. 2023; Chavarro-Carrero *et al*. 2024; Mesny *et al*. 2024). While these effectors exhibit antimicrobial activity, it remains unclear whether they additionally perform other functions, such as manipulation of host physiology. Using a gnotobiotic system, we now demonstrate that it is possible to address this question by testing virulence contributions separately, in the absence and in the presence of plant-associated microbiota. We show that the antimicrobial effector Ave1L2, which was previously shown to facilitate host colonization through suppression of Actinobacteria (Snelders *et al*. 2023), does not markedly contribute to virulence during infections on tomato plants that were grown in sterilized substrate. Notably, it contributes to virulence in the presence of a plant-associated microbiota, suggesting that Ave1L2 enhances fungal virulence through its antimicrobial activity, and furthermore that it lacks significant other virulence functions, and thus host targets. These findings contrast with those for the *V. dahliae* effector Ave1 that was previously shown to facilitate host colonization through suppression of Sphingomonad bacteria (Snelders et al. 2020), as we now reveal that Ave1 also significantly contributes to virulence on tomato plants grown under sterile conditions. This finding points towards a dual role for the Ave1 effector, contributing to microbiota manipulation as well as to host manipulation.

Multifunctionality has previously been observed for other fungal effectors as well. For instance, *Parastagonospora nodorum* secretes Snf1, which induces cell death to promote nutrient release while also protecting the fungus from wheat chitinases (Liu *et al*. 2016). Likewise, *Ustilago maydis* utilizes the effector UmPr-1La to suppress host immunity while simultaneously sensing plant-derived compounds to guide hyphal growth (Lin *et al*. 2023). The Ave1 effector has widespread plant homologs and was likely acquired via horizontal gene transfer from plants (de Jonge *et al*. 2012). Notably, most plant homologs of Ave1 are annotated as plant natriuretic peptides (PNPs), systemically mobile molecules released during biotic and abiotic stress that play key roles in plant growth and homeostasis (Gehring and Irving 2003; Ruzvidzo *et al*. 2011; de Jonge *et al*. 2012). In this study, we demonstrate that Ave1 induces stomatal opening in the tomato epidermis, confirming its PNP activity *in vitro*, whereas Ave1L2 lacks this activity. Consequently, we speculate that Ave1 may contribute to virulence *in planta* by manipulating plant physiology, possibly through PNP-activity, thereby promoting host colonization. This mode of action is reminiscent of a strategy employed by the biotrophic bacterial plant pathogen *Xanthomonas citri* subsp. *citri*, which exploits the PNP homologue XacPNP to alter host physiology to promote infection and bacterial proliferation (Nembaware *et al*. 2004; Gottig *et al*. 2008; Garavaglia *et al*. 2010b; Garavaglia *et al*. 2010a).

Several studies have indicated that PNPs play roles in host defense against invading pathogens (Breitenbach *et al*. 2014; Ficarra *et al*. 2018). For example, expression of the Arabidopsis *AtPNP-A* gene is induced upon infection with *Pseudomonas syringae* pv. *syringae*, and *AtPNP-A* deletion mutants exhibit increased susceptibility to the pathogen (Ficarra *et al*. 2018). Notably, previous studies also reported that AtPNP-A, similar to Ave1, possess an antibacterial activity against *Bacillus subtilis* (Snelders *et al*. 2020). Thus, PNPs may similarly display dual activities and exert, antimicrobial effects besides their PNP activity that could contribute to shaping the plant microbiota as well as to antagonizing invading pathogens (Snelders *et al*., 2020). Future studies investigating the direct impact of PNPs on host microbial communities and pathogen susceptibility will be crucial to validate this hypothesis.

To facilitate mechanistic research on plant-microbiota interactions, which includes research into plant-pathogen interactions, various gnotobiotic systems have been developed (Innerebner *et al*. 2011; Carlström *et al*. 2019; Kremer *et al*. 2021; Ma *et al*. 2022). Among these, peat-based systems, such as the Flowpot-system, have emerged as valuable tools due to their ability to mimic natural conditions by providing a soil-like substrate structure, shielding roots from light, and supplying organic carbon (Kremer *et al*. 2021; Ma *et al*. 2022). To leverage the advantages of the Flowpot-system for studying plant-microbiota-pathogen interactions, we report inoculation protocols that allow to inoculate plants with *V. dahliae* in sterile settings. While *Arabidopsis thaliana* and *Lotus japonicus* were previously shown to be compatible with the Flowpot-system (Kremer *et al*. 2021; Wippel *et al*. 2021), we now also successfully established tomato growth in this system and established *V. dahliae* infections across all three host species. Additionally, we supplemented our gnotobiotic system by assembling a collection of 133 unique bacterial strains from tomato plants that were commercially grown on potting soil, capturing a wide diversity of tomato-associated microbiota. By providing full genome sequences for over half of our bacterial collection, we offer a valuable resource for mechanistic investigations into the tripartite interaction of plants, their microbiota and *V. dahliae*.

Collectively, our findings reveal that antimicrobial effectors can serve dual functions for fungal virulence, both as antimicrobial agents but also at the same time as modulators of host physiology. Notably, many antimicrobials are ancient proteins, widely distributed across the fungal kingdom, and likely functioned in microbial competition long before the evolution of land plants (Snelders *et al*. 2021; Snelders *et al*. 2022; Mesny and Thomma 2024). It is therefore conceivable that some of these effectors have adapted novel functions throughout co-evolution with host plants, potentially contributing to plant manipulation. This suggests that dual functionality may be a common feature among antimicrobial effector proteins. Furthermore, the development of *V. dahliae* inoculations in a tomato-compatible Flowpot-system combined with a tomato-associated bacterial collection provides a robust platform for future research on the role and contribution of antimicrobial effector proteins for fungal pathogens. Understanding the precise molecular mechanisms underlying the role of antimicrobial effector proteins to the biology of fungal plant pathogens may ultimately open up novel strategies for microbiota-based disease control in agriculture.

## Materials and Methods

### Preparation and assembly of the Flowpot-system

Flowpot substrate was prepared by sieving potting soil (Balster Einheitserde, Frödenberg, Germany) through a 1 cm x 1 cm mesh, using only the material that passed through. Vermiculite with a kernel size ranging from 0.1 mm - 0.3 mm (LIMERA Gartenbauservice, Geldern-Walbeck, Germany) was sieved through a 1 mm x 1 mm mesh, retaining only the material left in the sieve. The two components were mixed in a 1:1 ratio, followed by the addition of 150 ml of water per liter of substrate and thorough mixing, and autoclaved on a liquid cycle at 121°C for 20 minutes (Systec, Linden, Deutschland). After 16 hours of incubation in darkness, the substrate was mixed thoroughly in a sterile hood before 50 ml of water per liter of substrate were added and the substrate was autoclaved again on a liquid cycle. To assemble individual Flowpot units, truncated (at the 45-ml mark) and autoclaved 50 ml luer-lock syringes (Terumo Europe, Leuven, Belgium) were filled with an autoclaved 250 µm pore-size polyamide filter mesh (Biologie-Bedarf Thorns, Deggendorf, Germany), and a 3 cm layer of autoclaved 3 mm silica glass beads (Roth, Karlsruhe, Germany). Assembled Flowpots containing sterile substrate were autoclaved again on a liquid cycle. For re-colonized substrate, sterile substrate was mixed with 10% non-autoclaved substrate and incubated overnight at room temperature, after which the mixture was used to assemble Flowpots. To remove toxic compounds released during autoclaving, the substrate was flushed using a vacuum system. To this end, Flowpots were placed onto a Qiavac 24 plus system with luer-lock adapters (Qiagen, Venlo, The Netherlands). Vaccuum was applied and 30 ml sterile MQ water was poured into each Flowpot. Subsequently, the substrate was enriched by flushing with 30 ml nutrient solution. For tomato and Arabidopsis plants, half-strength Murashige & Skoog (MS) medium (Duchefa, The Netherlands) was added whereas for Lotus plants 0.25x B&D (Broughton and Dilworth 1971) solution was added. For SynCom treatments, the SynCom was added to the medium prior to flushing.

To assess sterility of the substrate, 500 mg substrate were suspended in 10 ml 100 mM MgCl_2_ and shaken at 300 rpm for 1 hour at room temperature. Following 1.000-fold dilution, the samples were plated onto lysogenic broth agar (LB), tryptic soy agar (TSA) and Reasoner’s 2A agar (R2A) and incubated at RT for up to 4 days.

### Plant material and seed sterilization

Plants used in the Flowpot-system were tomato (*Solanum lycopersicum* L.) cultivar MoneyMaker, *Arabidopsis thaliana* Col-0 and *Lotus japonicus* Gifu. Tomato and Arabidopsis seeds were surface sterilized as described previously (Schlesier *et al*. 2003). Following sterilization, the seeds were stratified at 8℃ for 24 hours and then sown into each Flowpot unit. Lotus seeds were rubbed with sand paper and sterilized by 20 minutes incubation in 10 ml of MQ water with 200 µl NaCOl on a rotary shaker at 185 rpm. Subsequently, seeds were washed 5 times with sterile MQ water. Sterilized Lotus seeds were germinated on 0.8% plant agar at 22℃ for 5 days before seedlings were transferred into Flowpot units. In total, five individual Flowpot units were placed into a Microbox container with four air-filters (SacO2, Deinze, Belgium) and placed in a greenhouse chamber (17 hours of light at 23℃ followed by 7 hours of darkness at 22℃).

### Fungal inoculation assays

For *Verticillium dahliae*inoculations, conidiospores were harvested from wild-type or effector deletion strains of *V. dahliae* (Snelders *et al*., 2020; Snelders *et al*., 2023) after growth on potato dextrose agar (PDA; Carl Roth, Karlsruhe, Germany) for 10 days. Conidiospores were washed three times by centrifugation at 10.000 rpm for 10 minutes followed by pellet resuspension in sterile MQ water. Conidiospores were counted using a Neubauer Chamber and the concentration of the final conidiospore-suspension was adjusted to 10^6^ conidiospores/ml. Inoculation was performed on plants that were grown for 14 days in the Flowpot-system. To this end, microboxes with Flowpots were opened in a sterile hood and plants were carefully uprooted from the Flowpots. Roots were rinsed with sterile MQ water and subsequently placed into the *V. dahliae* conidiospore suspension for minimum 8 minutes. Subsequently, plants were placed back into the original Flowpots and the boxes were placed back into the greenhouse. Verticillium wilt symptom development was monitored at 14 days post inoculation. Symptoms were monitored by measuring shoot fresh weight and canopy areas were calculated from overhead pictures using ImageJ (Schneider *et al*. 2012).

### Colony picking-based collection of tomato-associated bacteria

To assemble a collection of tomato-associated bacteria, root and stem samples from two commercially purchased tomato plants grown in a potting soil, were separated and cut into 2 mm long pieces and washed in 100 mM MgCl_2_. Subsequently, 3 mm metal beads were added and samples were homogenized for 3x 45 seconds at 30 Hz in a tissue-lyzer (Retsch, Haan, Germany). Samples were diluted 1/10;1/100;1/1.000 and plated on agar plates containing either TSA, LBA, M9 minimal medium, R2A or R2A supplemented with 0.5% v/v. Plates were incubated in darkness at room temperature for 5 days and individual colonies were picked and transferred onto fresh plates. Following 2 rounds of single colony streaking, material of each colony was added to 1 ml of sterile 25% glycerol and stored at −80℃. To identify the bacteria, a 5 μl loop of bacteria was transferred into 1 ml of MgCl_2_ and vortexed for 5 minutes. DNA was extracted as described previously (Zhang *et al*. 2021) and used to amplify the 16s rRNA gene using the 27F (AGAGTTTGATCCTGGCTCAG) and 1492R (GGWTACCTTGTTACGACT) primers. PCR was conducted using Phusion HF polymerase (New England Biolabs, Ipswich, USA) at 98℃ for 30 s followed by 30 cycles of 98℃ for 10 s, 56℃ for 30 s, 72℃ for 45 s and a final extension at 72℃ for 8 min. PCR products were examined on 1.5% agarose gel and purified using the ExoSAP-IT Express Kit (Thermo Fisher Scientific, Waltham, USA). Following purification, the samples were collected in 96-well plates and send for Sanger sequencing (Microsynth Seqlab, Göttingen, Germany). Sequences were trimmed using CLC workbench (Qiagen, Venlo, The Netherlands) and blasted against the NCBI rRNA/ITS database. Identified strains were used to construct a phylogenetic tree of the collection with the ETE-Toolkit (V3.1.3, Huerta-Cepas *et al*. 2016). Visualization of the tree was conducted using iTOL (V6.9.1, Letunic and Bork 2024).

### SynCom preparation

To generate a disease-suppressive SynCom, all isolated Sphingomonadales strains from the collection of root-associated tomato bacteria were selected. Additionally, one representative strain from each other bacterial family in the bacterial culture collection was selected, with preference for plant-beneficial strains; if none were identified, a random strain was chosen. All selected bacterial strains were cultured in R2A broth for 2 days at room temperature and 160 rpm. Bacterial cells were collected by 10-minute centrifugation at 4500 rpm. Cells were washed twice with 10 mM MgSO_4_ and resuspended in half-strength MS to an OD_600_ of 0.5. All cultures were combined in equal amounts and the overall OD_600_ of the final SynCom was adjusted to 0.02. Next, the SynCom was applied to the Flowpot-system by flushing the soil.

### Tomato microbiota sequencing

Flowpot tomato plants were harvested in a sterile hood and ground to powder using a tissue lyzer (Retsch, Haan, Germany). DNA was extracted using the Power Soil Pro Kit (Qiagen, Venlo, The Netherlands). DNA was further purified using the Monarch PCR&DNA Clean up Kit (New England Biolabs, Ipswich, United States). All kit-solutions were filter-sterilized before use. Purified DNA was used to amplify the V3-V4 region of the 16S gene in the presence of the pPNA and mPNA blocking clamps (PNABio, Newbury Park, United States). Amplicons were sequenced using 16S sequencing on an Illumina MiSeq Platform (BGI-Genomics, Shenzhen, China). To sequence the input material for the culture collection, root and stem samples of the tomato plant were manually ground. Subsequently DNA was extracted as described previously (Chavarro-Carrero *et al*. 2021). Extracted DNA was used to amplify the V5-V7 region of the 16S gene using 799F and 1193 primers as described previously (Wippel *et al*. 2021). Purified amplicons were submitted for sequencing on an Illumina sequencing platform (Cologne-Center for Genomics, Cologne, Germany). Data analysis was conducted as described previously (Callahan *et al*. 2016; Snelders *et al*. 2020).

### Nanopore Sequencing and bacterial genome assembly

Bacteria were cultured in R2A-broth for 48 hours and pelleted through centrifugation. Bacterial pellets were resuspended in TEN-Buffer (10 mM Tris-HCL pH 8.0, 10 mM EDTA, 150 mM NaCl), supplemented with 20 µl lysozyme (20 mg/µl) and incubated at 37℃ for 20 minutes. Next, 3 µl RNase A (20 mg/µl) were added and the samples were incubated at 65℃ for 5 minutes. Subsequently, 550 µl of a reduced TEN-buffer (10 mM Tris/HCL, 1 mM EDTA, 50 nM NaCl), supplemented with 2 µl of proteinase K (20 mg/µl) and 40 µl SDS (10% w/v) were added followed by incubation for 2 hours at 60℃. Subsequently, phenol washing was performed twice and the aqueous phase was further cleaned by two chloroform washing steps. Next, DNA was precipitated by adding 10 volumes of ice-cold 100% EtOH and incubation at 4℃ overnight. Precipitated DNA was collected and washed with 70% EtOH and resuspended in MQ water. DNA quality and quantity were assessed using Qubit, Nanodrop and agarose gel assays. Full genome sequencing was carried out on a Nanopore MinION device using R10 Flowcells (Oxford Nanopore Technologies, Oxford, UK). The sequencing library was prepared using the ligation sequencing gDNA-Native Barcoding Kit 96 V14 (SQK-NBD114.96; Oxford Nanopore Technologies, Oxford, UK). Bacterial genomes were assembled using the uncorrected sequenced reads in Flye (2.9.5) with default settings and the --nano-hq input option (Kolmogorov *et al*. 2019). The assembled genomes were annotated using Prokka (1.14.6) and completeness of the genome assemblies was assessed with BUSCO (5.3.2) (Manni *et al*. 2021; Seemann 2014).

### Protein production and stomatal opening assay

Protein sequence alignment was performed using MAFFT (Version 7.271; Katoh *et al*. 2002). The sequences encoding mature Ave1 and XacPNP were cloned into the pET-15b expression vector for N-terminal His_6_ tagging (Novagen, Madison, WI, USA) (for primer sequences see Supplementary Table 1). Heterologous proteins were produced as described previously (Snelders *et al*., 2020) and purified from inclusion bodies under denaturing conditions using His60 Ni^2+^ Superflow Resin (Clontech, Mountain View, CA, USA). Purified proteins were stored in 0.25 M ammonium sulphate with 0.1 M BisTris, pH 5.5. Final concentrations were determined using the BioRad Protein Assay (BioRad, Veenendaal, The Netherlands). Stomatal aperture was tested as described previously (Gottig *et al*. 2008) using tomato leaf tissue.

## Author contributions

W.P., K.W. and B.P.H.J.T. conceived the project. W.P., H.R., K.W. and B.P.H.J.T. designed the experiments. W.P., J.P., A.K., N.S., J.W., N.C.S., G.L.F., E.A.C.-C., A.L.M., G.C.P. and N.S. performed the experiments. W.P., A.K., H.R. and B.P.H.J.T. analyzed the data. W.P. and B.P.H.J.T. wrote the manuscript. All authors read and approved the final manuscript.

## Acknowledgements

A.L.M. is holder of a postdoctoral research fellow funded by the ‘Fundación Ramón Areces’. K.W. received funding from the German Research Foundation (DFG) special priority programme Deconstruction and Reconstruction of the Plant Microbiota (DECRyPT SPP2125, project 466384394) and from the Research Priority Area Systems Biology of the Faculty of Science at University of Amsterdam. B.P.H.J.T. acknowledges funding by the Alexander von Humboldt Foundation in the framework of an Alexander von Humboldt Professorship endowed by the German Federal Ministry of Education and is furthermore supported by the Deutsche Forschungsgemeinschaft (DFG, German Research Foundation) under Germany’s Excellence Strategy – EXC 2048/1 – Project ID: 390686111.

## Data availability

Genomes from the bacterial culture collection are available via the European Nucleotide Archive (ENA) under the study accession number PRJEB87606. The genome annotation outputs as well as the scripts used in this study are available on https://github.com/antonkraege/BacterialTomatoCultureColection_Genomes/

