## Supplementary Figures for "A gnotobiotic system reveals multifunctional effector roles in plant-fungal pathogen dynamics"

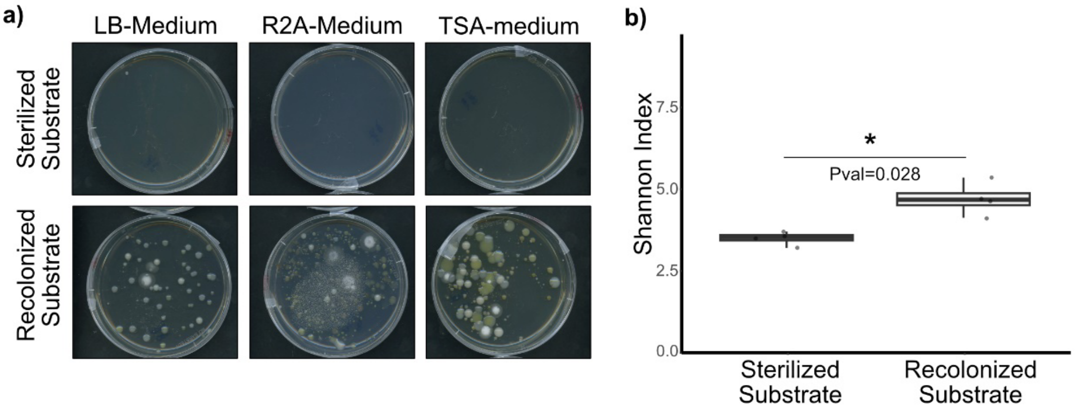


**Supplementary Figure 1 a)** Microbial growth from sterilized or recolonized Flowpot substrate on three growth media after four days of incubation. **b)** Bacterial alpha diversity in stem tissue of tomato plants grown on sterilized or recolonized Flowpot substrate. Pairwise test using Wilcoxon rank sum test (Pval < 0.05).

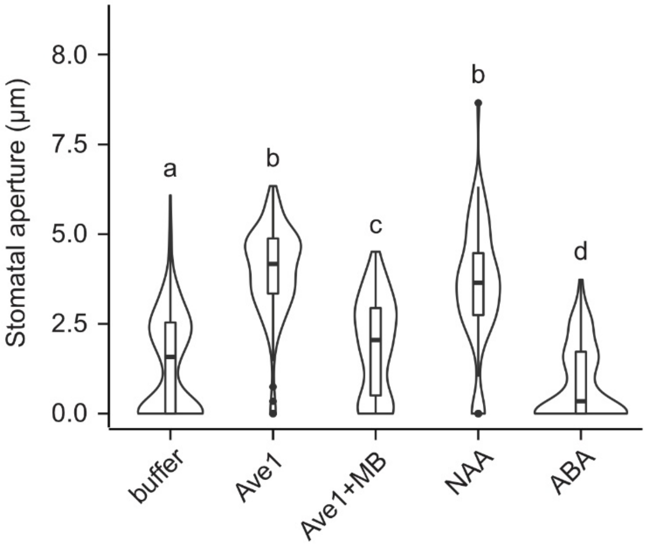


**Supplementary Figure 2. PNP activity of VdAve1 is mediated by cGMP signaling.** Stomatal opening in tomato epidermis following treatment with 5 µM VdAve1 with or without the cGMP signaling inhibitor methylene blue (MB). Indole-3-acetic acid (IAA; 1 µM) and 50 µM abscisic acid (ABA) were used as positive and negative controls, respectively. Data are from one representative experiment. Experiments were performed twice. Letters represent statistically significant differences in stomatal opening according to one-way ANOVA ($F (5,824) = 124.8$, $p<0.001$) and Tukey's post-hoc test. Error bars represent the mean ± SD ($n>70$).
